# Optimizing gene expression by adapting splicing

**DOI:** 10.1101/220228

**Authors:** Idan Frumkin, Ido Yofe, Raz Bar-Ziv, Yoav Voichek, Yitzhak Pilpel

## Abstract

Can splicing be used by cells to adapt to new environmental challenges? While various adaptation mechanisms for regulating gene expression have been revealed for transcription and translation, the role of splicing and how it evolves to optimize gene-expression patterns has not been thoroughly investigated. To tackle this question, we employed a lab-evolution experimental approach that challenged yeast cells to increase expression levels of a gene that carries an inefficiently-spliced intron. We followed the evolution of multiple lines and found independent routes by which cells adapted. Surprisingly, we did not observe an intron loss event, a mechanism believed to be common in intron evolution. Instead, we identified mutations in *cis* that improved the intron’s splicing efficiency and increased the overall expression level of the entire gene. One of these *cis*-acting mutations occurred in an adjacent exon and hampered the functionality of the gene that was not under selection - demonstrating that adaptation of splicing efficiency may sometimes come at the expense of protein activity. Additionally, we observed adaptations in *trans*, which increased the cellular availability of the splicing machinery. These adaptations were achieved either by elevated expression levels of the splicing apparatus or, unexpectedly, by reduced expression levels of other intron-containing genes that are the natural consumers of this process. Ultimately, our work reveals novel molecular means by which the splicing machinery is changed by natural selection to optimize gene-expression patterns of cells.

## Introduction

Changing environments can force cells to change their gene-expression programs, to better accommodate their surroundings. Throughout evolution, cells acquired regulatory mechanisms to tune gene expression, which have been the subject of intensive investigations – focusing mainly on transcription and translation. For example, when cells are challenged to increase protein expression levels, the DNA sequence of genes can change so as to increase transcription^1,2^, support more efficient mRNA translation^3,4^, or result in greater mRNA transcript stability^5,6^. Additionally, the transcription and translation machineries themselves have been shown to adapt to environmental challenges by altering the cellular pools of transcription factors^7^ or tRNAs^8,9^.

In evolving expression programs, adaptation often occurs either directly on the genes under pressure (“evolution in *cis*”)^10^, or indirectly, *e.g.* on the expression machineries, mostly transcription and translation (“evolution in *trans*”)^11,12^. These two routes of evolution are profoundly different, as the first (*cis*) provides a localized solution that in principle can affect only a certain gene, while the later (*trans*) could be the method of choice if a coordinated change in many genes is needed.

Surprisingly, although the process of splicing is central to the maturation and regulation of mRNAs in eukaryotes^13–17^, its role in adapting to novel demands on gene expression has not been thoroughly investigated. During mRNA splicing, precursor mRNAs are processed to remove introns while fusing exons together to create the mature transcript. This process provides an evolutionary means to diversify the proteome towards phenotypic novelty, as the choice of intron to be excluded, as well as the exons which are found in the mature transcript, can both be regulated based on the cell’s needs^15,18,19^. One aspect of splicing evolution that has been extensively studied is gain and loss of intronic DNA, for which several molecular models have been proposed, mainly Reverse-Transcription and recombination-mediated intron loss, intron transposition and also exonization and intronization via mutations^20–23^. While intron loss and gain have been demonstrated experimentally^24,25^, other forms of splicing evolution, such as alterations in splicing efficiency under changing conditions, have not.

Here, we set out to reveal whether introns or the splicing apparatus can evolve so as to alter the expression levels of genes in a timely and adaptive manner, and ask whether and how splicing evolves in *cis* and in *trans* to regulate gene expression. To this end, we generated a reporter construct in yeast cells that could simultaneously be read out and be selected for splicing efficiency. Namely, we introduced an inefficiently-spliced intron to a reporter gene that was fused to an antibiotic resistance gene. Using this approach, we could carry out a lab-evolution experimental setup to study the adaptation of splicing in the presence of the corresponding antibiotics.

Our results demonstrate two alternative adaptive routes for this evolutionary challenge. First, *cis*-acting solutions, in the form of adaptive mutations, occurred in the intron itself, but also surprisingly in an up-stream exon. These mutations resulted in increased splicing efficiency and higher expression levels of the antibiotic resistance gene. Remarkably, in some evolutionary lines there were no *cis* mutations, but rather *trans*-acting adaptations that have increased cellular availability of the splicing machinery - either by increased expression levels of the machinery’s genes themselves, or surprisingly, by decreasing the expression levels of other intron-containing genes. Thus, this works unravels different layers at which the splicing machinery can be adapted to alter gene expression.

## Results

### Low splicing efficiency leads to stressed cells under restrictive conditions

We hypothesized that tuning splicing of genes could serve as a means to optimize their expression levels. To test this hypothesis, we used the yeast *Saccharomyces cerevisiae* in which ˜30% of the transcriptome must be spliced, at a range of splicing efficiencies^17,26^, to form mature mRNAs^27^. We built a synthetic gene construct that consists of two fused domains: A fluorescent reporter (YFP), which includes two alternative natural introns - with either high or low splicing efficiency - near the YFP’s fluorescence site^26^, fused to an antibiotics resistance gene (Kanamycin resistance gene). Specifically, we created three strains: (i) WT YFP strain without an intron; (ii) “Splicing^High^” in which the YFP harbors the natural intron of *OSH7* and was previously reported to have high splicing efficiency within this YFP context^26^; and (iii) “Splicing^Low^” in which the natural intron of *RPS26B,* with a low splicing efficiency^26^, was inserted in the same location (Figure 1A).

**Figure 1.**
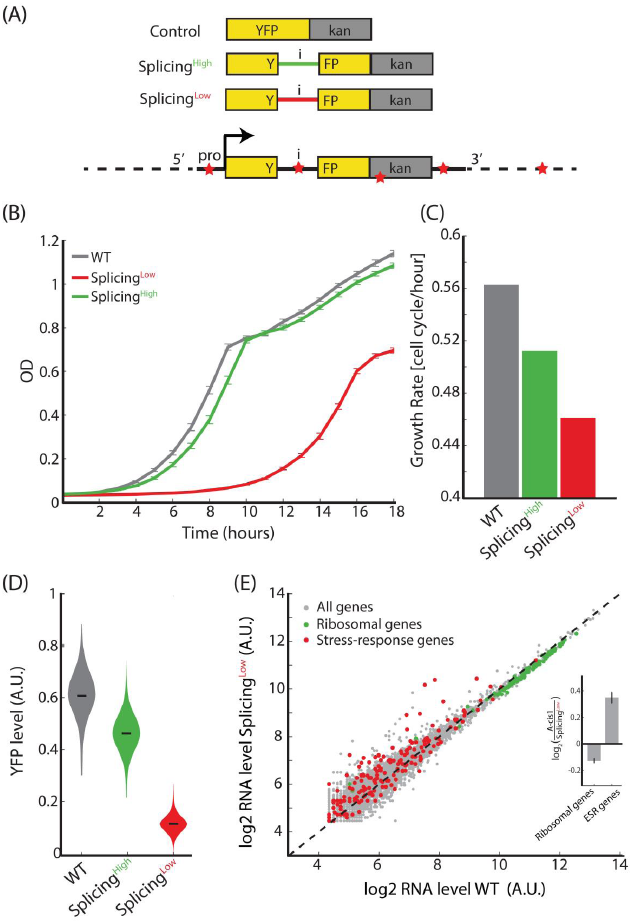
– Inefficient intron splicing leads to lower gene expressionlevelsandcompromisedantibiotics resistance. **A|** We introduced two alternative introns into a YFP domain that was fused to a kanamycin resistance domain - to generate three strains: (i) WT without an intron; (2) Splicing^High^ with an efficiently spliced intron; and (iii) Splicing^Low^ with an inefficiently spliced intron. Evolving cells at the presence of the antibiotics could adapt by mutating different parts of the YFP-Kan construct (evolution in cis) or other loci, evolution in *trans* (red stars represent potential locations of such putative mutation sites). **B+C|** Splicing^Low^ suffers from a severe growth defect compared to WT or Splicing^High^ cells when the antibiotic is supplemented to the medium. The growth defect is manifested as a longer lag phase and a lower maximal growth rate. **D|** Florescence intensity of the YFP-Kan reporter for all three strains shows that Splicing^Low^ cells have lower expression levels of YFP-Kan. This observation links between YFP-Kan expression levels and cellular fitness. **E|** Transcriptome profiling shows that ribosomal genes were down-regulated (green dots, p-Value=4.62x10^−26^, paired t-test) and stress-related genes were up-regulated (red dots, p-Value=3.40x10^−5^, paired t-test) in Splicing^Low^ compared to WT cells. This observation suggests that Splicing^Low^ cells are stressed because of compromised resistance to the antibiotics and that the general stress response was activated in them. **Inset|** Mean log_2_ ratio of ribosomal and ESR gene groups.

We first hypothesized that cellular growth of each strain in the presence of the antibiotics, geneticin (G418), will associate with the YFP-Kan expression levels. We followed the growth of the three strains in the presence of the antibiotics and found that the WT strain had the highest fitness, Splicing^High^ grew slower, and Splicing^Low^ demonstrated a severe growth defect compared to the two other strains (Figure 1B+C). We then measured florescence intensity of the YFP-Kan reporter in the presence of the drug. In line with the growth measurements, we observed that WT cells demonstrated the highest fluorescence levels, followed by Splicing^High^, and with Splicing^Low^ cells showing the lowest YFP-Kan levels (Figure 1D). These results demonstrate that the inefficiently-spliced intron in Splicing^Low^ reduces cellular levels of YFP-Kan and hence lead to a reduced fitness.

Since YFP-Kan expression level in Splicing^Low^ were significantly lower compared to the other strains, we hypothesized that Splicing^Low^ cells did not reach the needed concentration to sufficiently neutralize the antibiotics, and hence resulted in stressed cells. To test this hypothesis, we performed mRNA sequencing of exponentially growing WT and Splicing^Low^ cells in an antibiotics containing medium, and analyzed their transcriptome profiles. Indeed, we observed that ribosomal genes were down-regulated in Splicing^Low^ compared to the control strain – a clear signature of stressed cells^28^ (Figure 1E). Notably, the reduction in ribosomal expression levels (˜8%) we observed here due to growth rate differences between WT and Splicing^Low^ cells is accurately predicted by a recent study, which calculated the linear correlation between growth rate and ribosomal expression levels in yeast cells^29^. In parallel, stress-related genes^30^ were up-regulated in the Splicing^Low^ compared to the control strain (Figure 1E). We thus concluded that the general stress response was activated in Splicing^Low^ cells.

### Rapid evolutionary adaptation increases expression level of the resistance gene

Our experimental system mimics an evolutionary scenario in which there is an immediate and continuous selection pressure to up-regulate the expression level of a gene of interest. How would the system evolve to better resist the antibiotics? Possible means to adapt include mutations in the gene’s promoter to increase transcription, mutations that increase translation initiation, or mutations inside the gene itself that increase the functional efficiency of the protein. Additionally, the splicing machinery may also take part in adaptation of gene expression levels. To find which evolutionary track would be used by cells, we evolved the three strains by daily serial dilution on a medium supplemented with G418 for ˜560 generations, in four independent cultures for each strain (Figure 2A). Interestingly, only the cultures of Splicing^Low^ cells demonstrated a significant improvement in fitness at the end of the experiment (Figure 2B+C). This observation implies that only Splicing^Low^ experienced a sufficiently strong selective pressure to adapt to the presence of the antibiotics in the medium, in contrast to the WT and Splicing^High^ strains which originally had much higher levels of the resistance gene.

**Figure 2.**
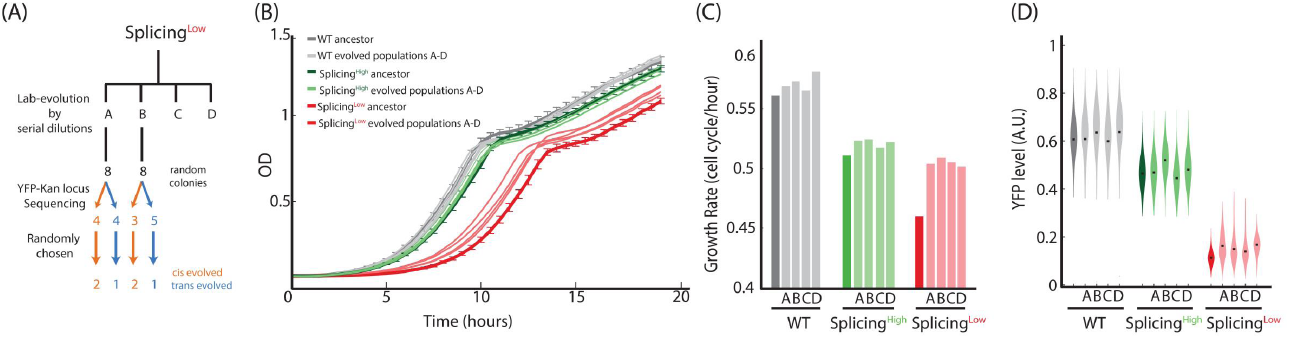
– Rapid adaptation to the presence of the antibiotics is observed only for Splicing^Low^ cells. **A|** We evolved WT, Splicing^High^, and Splicing^Low^ cells for ˜560 generations with the presence of the antibiotics in four independent cultures for each strain. We measured fitness and YFP-Kan expression levels for all evolved lines (see below), and also randomly chose 16 colonies from two evolved lines of Splicing^Low^. We sequenced the YFP-Kan locus of those colonies and observed that around half showed mutations in the YFP-Kan construct (indication of evolution in *cis*) and the other half did not (indication of evolution in *trans*). Of those colonies, we randomly chose two *cis*-evolved and one *trans*-evolved colonies from each evolved population for further examinations (see figure 3 onwards). **B+C|** Growth of evolved populations compared to the three ancestors. Only evolved Splicing^Low^ cells demonstrate significant improvement in growth for all four independent evolution lines. This observation suggests that the inefficiently spliced intron led to a rapid adaptation of Splicing^Low^ cells. **D|** Florescence intensity of the YFP-Kan reporter for all evolved cultures show that expression levels were much increased in all four evolved cultures of Splicing^Low^ compared to the ancestral strain (effect sizes = 78.67, 79.54, 75.17, 83.19). Conversely, the increase in expression levels in the evolved WT and of Splicing^High^ populations were smaller(WT effect sizes= 64.66,68.44,63.51,67.74; Splicing^High^ effect sizes=54.33, 70.66, 52.43 and 58.27). This observation suggests that adaptation of Splicing^Low^ cells was based on their ability to increase expression levels of the resistance proteins.

Consistent with the fitness measurements, YFP measurements of the evolved cultures showed that expression levels of the resistance-YFP fusion gene increased in all four evolved cultures of Splicing^Low^ compared to the ancestral strain (Figure 2D). Conversely, the increase in YFP-Kan expression levels in the evolved WT populations was smaller, and only one culture of the evolved Splicing^High^ cells demonstrated strong elevation of the YFP-Kan levels (Figure 2D). These results further support that Splicing^Low^ cells experienced the strongest selective pressure to adapt rapidly to the presence of the antibiotics in our experimental setup, and that they achieved this goal by increasing the levels of the resistance gene. We next moved to reveal the molecular mechanisms underlying this evolutionary process.

### Adaptation in *cis* and *trans* leads to increased splicing efficiency

We hypothesized that improving the low splicing efficiency of the intron in Splicing^Low^ could be exploited by natural selection as an adaptation mechanism towards increasing the resistance gene levels. We therefore sequenced the YFP-Kan locus in 16 randomly chosen colonies from two evolved populations (termed here population A and population B) of Splicing^Low^. Interestingly, we found that the colonies were split into two types – either with or without a mutation in the YFP-Kan locus. In population A, we found that the same mutation occurred in four out of eight colonies, changing adenine to cytosine inside the intron, 97 nucleotides up-stream to its 3’ end (Figure 3A). In population B, we identified an exonic non-synonymous mutation that changed a valine at position 61 of the YFP protein into alanine (a thymine to cytosine 14 nucleotides up-stream of the intron) in three out of eight colonies. In the five other colonies from this population there were no mutations in the YFP-Kan locus.

**Figure 3.**
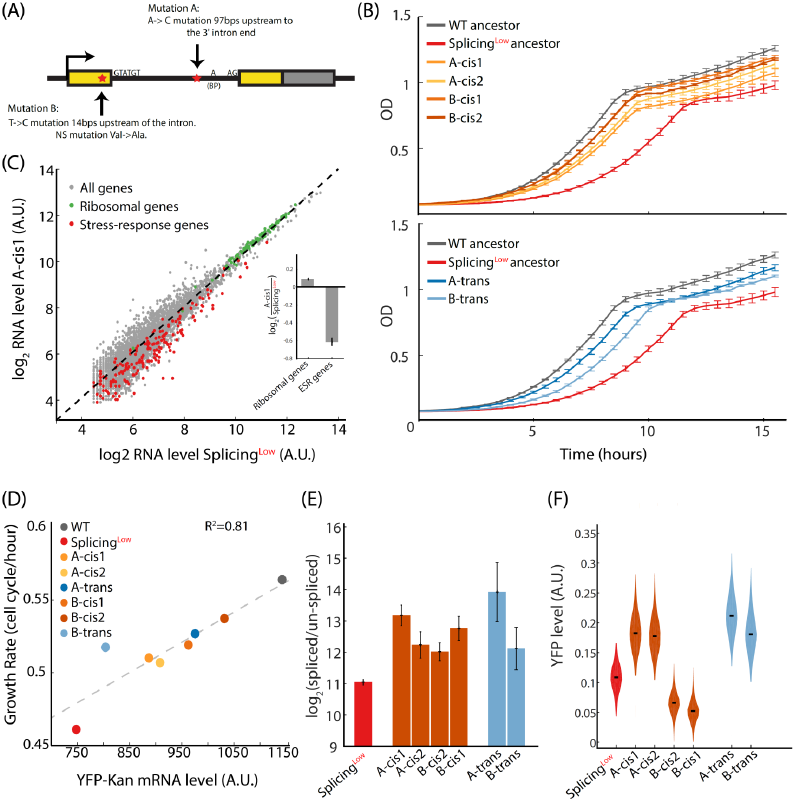
– Evolved colonies demonstrate increased splicing efficiency that results in higher transcript levels and relieved stress. **A|** Sequencing of the YFP-Kan construct in the evolved colonies revealed two mutation types: (i) in the intron itself and (ii) in the up-stream exon – see text for full description. These mutations did not occur in the intron 5’ donor, 3’ acceptor, or the branching point – suggesting that other positions of the intron and its vicinity are phenotypically functional and may affect splicing efficiency. **B|** All *cis*-evolved colonies (upper graph) and *trans*-evolved colonies (lower graph) show increased fitness compared to the Splicing^Low^ ancestor, yet still lower than the WT ancestor. **C|** Transcriptome profiling reveals that ribosomal genes were up-regulated (green dots, p-Value=4.94 x10^−18^, paired t-test) and stress-related genes were down-regulated (red dots, p-Value=3.64 x10^−15^, paired t-test) in the evolved colony A-cis1 compared to the Splicing^Low^ ancestor. This trend was observed in 5 out of 6 evolved colonies (Supplementary Figure 1). **Inset|** Mean log_2_ ratio of ribosomal and ESR gene groups. **D|** mRNA levels of YFP-Kan transcripts correlate with growth rate – suggesting that cellular fitness in our set-up is indeed determined by the availability of Kanamycin-resistance proteins to overcome the antibiotics. **E|** All *cis*- and *trans*-evolved colonies demonstrate increased splicing efficiency of the YFP-Kan mRNA compared to the Splicing^Low^ ancestor. This result suggests that all adaptation trajectories led to the adaptation of the splicing process to better mature the un-spliced YFP-Kan transcript. **F|** Florescence intensity of the YFP-Kan reporter show increased levels for the two *cis*-evolved colonies with the mutation in the intron and for the two *trans*-evolved colonies. In contrast, the two *cis*-evolved colonies with the non-synonymous mutation in the exon demonstrate decreased YFP-Kan levels. This observation suggests that the non-synonymous mutation hampered the ability of the YFP domain to florescent and reduced the Florescence intensity per protein molecule (see text for full explanation).

Notably, none of the colonies demonstrated a mutation in the construct’s promoter, terminator or in the sequence of the Kan resistance gene itself. These results propose that different mutations in the intron, or its vicinity, were adaptive and might affect splicing efficiency of the intron. Surprisingly, the observed mutations did not occur in the 5’ donor, 3’ acceptor, nor in the intron branch point – suggesting that other position of the intron can also be selected in evolution increase fitness by affecting splicing. While the intron- and exon-mutated colonies represent an evolutionary adaptation in *cis*, the colonies that showed no mutation in the entire gene construct potentially found adaptive solutions in *trans* that may have occurred elsewhere in the genome.

We randomly chose six colonies: four colonies with a *cis* mutation and two colonies that showed no mutations in *cis*, for which we reasoned that such colonies may have adapted in *trans*. We termed these colonies according to the evolution lines from which they were derived: A-cis1, A-cis2, B-cis1, B-cis2, A-trans and B-trans. We followed the growth of these evolved colonies in the presence of G418 and found, as expected, that all grew faster than the Splicing^Low^ ancestor (Figure 3B). We then performed RNA-seq and transcriptome analysis of all colonies, which revealed relaxation of the stress response that was featured in the ancestor. Namely, the general stress response genes were reduced and ribosomal proteins were up- regulated in five evolved colonies (Figure 3C and Supplementary Figure 1). These observations suggest that the cells indeed adapted to the presence of the antibiotics in the environment and that the stress experienced by them was partially alleviated.

We next hypothesized that cellular fitness might correlate with mRNA levels of the YFP-Kan construct because increased transcript levels should result in higher concentrations of the Kan protein. Indeed, maximal growth rates of the control and Splicing^Low^ ancestors and for the six evolved colonies correlate with mRNA levels of the YFP-Kan construct, as deduced from the RNA-seq (Figure 3D) – supporting our conclusion that adaptation was based on increasing expression levels of the YFP-Kan gene. Since the observed *cis* mutations occurred at the vicinity of the intron, we hypothesized that they increased splicing efficiency of the YFP-Kan transcript. To test this possibility, we performed, for both *cis*- and *trans*-evolved colonies, a splicing efficiency assay with qPCR - targeting the un-spliced and spliced transcript versions. Interestingly, the ratio of spliced to un-spliced transcripts was higher in all evolved colonies compared to the Splicing^Low^ ancestor, suggesting that at least some of the mRNA level increase we observed in the evolved colonies results from increased splicing efficiency (Figure 3E).

To prove that adaptation of the colonies actually led to higher protein levels of the resistance gene, we measured fluorescence intensity using flow cytometry. We found that the two *cis*-colonies from population A (A-cis1 & A-cis2) and the two *trans*-colonies (A-trans & B-trans) showed higher YFP-Kan levels compared to the ancestor. However, the two *cis*-colonies from population B (B-cis1 & B-cis2) demonstrated decreased fluorescence intensity values (Figure 3F). These observations suggest that the non-synonymous, exon mutation reduced the fluorescence-per-protein value of the YFP-Kan construct in these colonies. Indeed, this position corresponds to a position that was recently reported to reduce florescence when mutated in the highly similar GFP^31^. Because YFP functionality was not selected for or against in our setup, it was free to mutate as long as it helps achieve a higher expression level of the entire construct by increasing the intron’s splicing efficiency. It thus seems that modular domain-architecture of a protein may increase its evolvability under relevant conditions as it allows the optimization of each domain in isolation from the other.

It is possible that additional beneficial mutations exist in the genome of the *cis*-evolved colonies, which account for the phenotypes we observed. To directly assess the effects of the *cis* mutations, we generated two rescue strains, termed rescue-A and rescue-B, in which these *cis*-acting mutations were introduced individually to the ancestral Splicing^Low^ background. Notably, the two rescue strains grew better than Splicing^Low^ cells in the presence of the antibiotics (Figure 4A), though not as good as the wild-type, and the stress experienced by the Splicing^Low^ cells was relieved upon insertion of each individual *cis* mutation (Figure 4B). Finally, we measured splicing efficiencies and fluorescence intensity levels for both rescue strains, and found that they resembled the results of the evolved single colonies (Figure 4C-D, in comparison to Figure 3E-F). These observations strengthen our conclusions that the *cis*-acting mutations are sufficient to elevate YFP-Kan levels through an increased splicing efficiency, yet the non-synonymous mutation of population D also hampers the function of the YFP domain and reduces its florescence-per-protein ratio.

**Figure 4.**
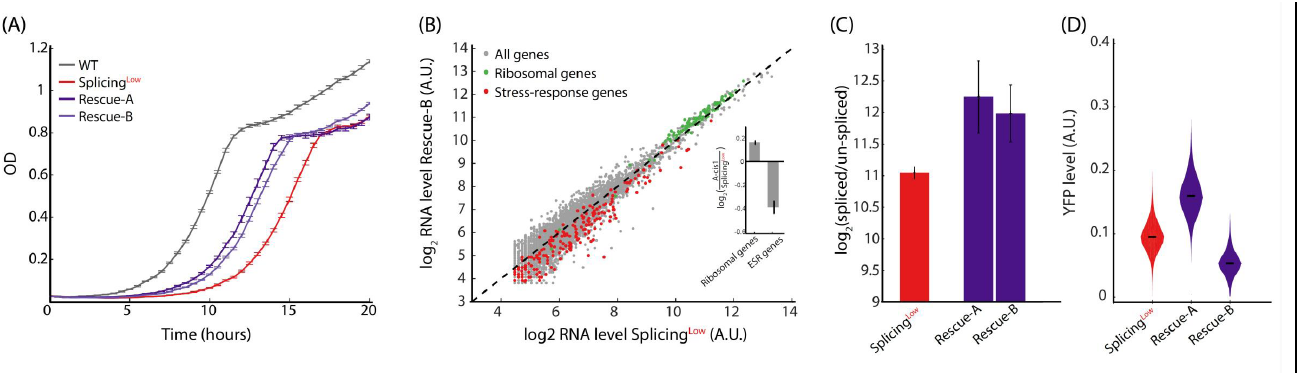
– cis-acting mutations are sufficient to increase fitness by elevating splicing efficiency. **A|** We created two rescue strains, each harboring one of the mutations that appeared spontaneously in the evolved populations. Growth of the two rescue strains show that a single mutation in the YFP-Kan construct is sufficient to increase fitness compared to Splicing^Low^. **B|** The exonic mutation is also sufficient to alleviate stress, as ribosomal genes were up-regulated (green dots, p-Value=1.02x10^−18^, paired t-test) and stress-related genes were down-regulated (red dots, p-Value=9.02x10^−12^, paired t-test) in Rescue-B compared to Splicing^Low^. The same trend was also observed for the intronic mutation for Rescue-A cells. **Inset|** Mean log_2_ ratio of ribosomal and ESR gene groups. **C|** The two rescue strains demonstrate higher splicing efficiency of the YFP-Kan mRNA compared to the Splicing^Low^ ancestor. This result suggests that a single mutation is sufficient to improve splicing efficiency. **D|** Florescence intensity of the YFP-Kan reporter for the Rescue-A and Rescue-B strains show similar trends as the colonies in Figure 3D - supporting earlier conclusions.

Our results thus far provide direct evidence that intron splicing takes part in the adaptation and optimization of gene expression patterns to environmental needs. Although intron sequences are much less conserved compared to exons, and are believed to be less functional, we demonstrate that their sequence can be used by natural selection as a molecular mechanism to regulate splicing efficiency and adjust gene expression patterns.

### Increasing cellular availability of the splicing machinery can be adaptive

We finally aimed to decipher the mechanism behind the increased YFP-Kan levels in the *trans*-evolved colonies that showed no mutations in *cis*, i.e. within the reporter gene or in its vicinity. We reasoned that elevating availability of the splicing machinery as a global resource could be a means to increase splicing efficiency of the YFP-Kan transcript, and thus could be used as an adaptive mechanism to the antibiotics challenge. Increased splicing-availability could be achieved by increasing the expression of the splicing machinery genes. In addition, as with other cellular machineries whose functioning depends on supply-to-demand economy^4,8,32,33^ reducing expression levels of the intron-containing genes, namely the “demand”, could increase the availability of the machinery towards the intron under selection here.

To test if any of these evolutionary routes were indeed taken by the evolved cells, we calculated the expression level ratio of genes between the evolved colonies and their ancestor. In colony A-trans, we observed that while the average expression-ratio of the splicing machinery genes (the “supply”) increased, that of the non-ribosomal intron-containing genes (the “demand”) decreased (Figure 5A). This observation suggests that indeed the cellular availability of the splicing machinery was elevated in this evolved colony, which might have allowed for the observed increased splicing efficiency of the YFP-Kan gene. Next, we hypothesized that the *cis*-evolving colonies may have also adapted in *trans* and used this adaptation mechanism as well. Indeed, in all other evolved colonies we observed a similar trend, in which the overall supply-to-demand measurement of the splicing machinery was increased (Figure 5B). Notably, in some colonies this phenotype was achieved by only increasing expression levels of splicing genes or by only reducing levels of the intron-containing genes (Supplementary Figure 2). Importantly, the two rescue strains, which did not evolve and only harbor our artificially introduced *cis*-acting mutation, did not show any change in splicing availability (Figure 5B), strongly supporting our conclusion that this phenotype was achieved by further adaptation of the cells during our lab-evolution experiment. Thus, we concluded that both *cis* and *trans* adaptation routes can co-occur in the same genome towards optimization of its gene expression patterns.

**Figure 5.**
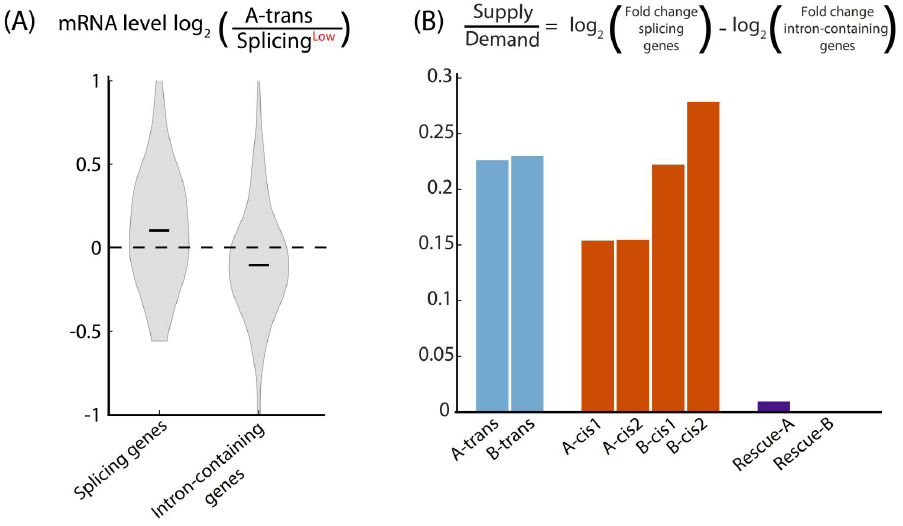
– Increasing cellular availability of the splicing machinery is an adaptive mechanism of splicing. **A|** The groups of splicing genes and intron-containing genes were increased (p-Value=1.36x10^−3^, paired t-test) and decreased (p-Value=1.67x10^−2^, paired t-test), respectively, in the trans-evolved colony A-trans compared to Splicing^Low^ ancestor. This observation suggests that the supply-to-demand ratio of the splicing machinery was increased in A-trans colony, which allowed its increased splicing efficiency of the YFP-Kan transcript. **B|** Supply-to-demand ratios for the splicing machinery were calculated to all evolved colonies and to the rescue strains as the difference between the mean fold-change of splicing genes to the mean fold-change of intron-containing genes. While supply-to-demand ratios were increased in all evolved colonies, they remained the same for the two rescue strains. These results suggest that indeed the cellular availability of the splicing machinery was elevated in the evolved colonies – a *trans*-adaptation mechanism to optimize gene expression using the splicing process.

## Discussion

Here we study the role of the splicing machinery in optimization of gene expression programs by placing selective pressure on cells to improve the splicing efficiency of a specific gene. Our results provide molecular evidence for the relevance of splicing as another instrument in the cellular toolbox towards adjusting its gene expression patterns. To the best of our knowledge, we demonstrate the first experimental evidence of splicing efficiency adaptation, confirming that this adaptation can occur in *cis* and *trans* similarly to adaptations of other means of gene regulation.

Two potential solutions to the burden we imposed on our ancestor lines were, surprisingly, not realized during our lab-evolution. First, considering previous studies of splicing evolution, one could have expected the intron to be lost by a genomic deletion or reverse transcription^34,35^. Such a solution could have been an ideal evolutionary adaptation to alleviate the burden, as we show that the intron-less strain has the highest fitness. The fact that we did not observe an intron-loss event suggests that nucleotide substitutions were more accessible solutions in this case, in agreement with previous evidence in yeast that nucleotide mutations are much more prevalent than deletion events, at a ratio of 33:1^36^. Another surprise was that the mutations we observed did not occur within any of the intron’s three functional sites: the 5’ donor, 3’ acceptor, or the branch point of splicing. Surprisingly, one mutation was actually detected, and was verified here to affect splicing, in a region of the intron not known to exert a major effect on splicing, and another splicing-improving mutation happened in the up-stream exon - suggesting that various positions in the intron and its proximity may facilitate splicing rate and take part in the evolution of this process.

Notably, the fluorescence intensity per protein molecule of the YFP domain was decreased due to the non-synonymous mutation in the YFP first exon, suggesting that under certain evolutionary constrains selection may hamper superfluous functions of certain protein domains so as to increase availability of the entire protein. Why then, would the mutations we observed increase splicing efficiency? Past evidence showed that mRNA secondary structures at the intron’s edges influence splicing efficiency^26,37–39^. It is possible then that the mutations we observed in this study somehow open the structure of the intron under selection and make it more accessible for the splicing machinery.

Adaptive changes also occurred in *trans* to the YFP-Kan locus and increased availability of the splicing machinery. Recently, the competition of pre-mRNAs for the splicing machinery was shown to affect cellular function, as splicing efficiency of multiple introns was influenced by changes in the composition of the transcript pool^40^. While this mechanism was elegantly suggested to maintain the separation between meiotic and vegetative gene expression states, it is also possible that it can be utilized as an adaptive route available for cells to optimize expression levels of genes. More broadly, it has been shown^41^ that yeast species that have a high content of intron-containing genes have adapted the codon usage of their splicing machinery genes more vigorously to their cellular tRNA pools compared to other species that have a lower number of introns in the genome. This evolutionary trend indeed shows that the splicing machinery adaptively responds to meet its own evolutionary demand.

Our findings demonstrate how availability of the splicing apparatus may have been beneficially increased both by elevating the expression level of the machinery and/or by reducing other intron-containing genes that compete with the antibiotic resistance un-spliced RNA for the spliceosome. Thus, increase in supply-to-demand ratio, analogous to the case in translation systems^8,42^, appears to have evolved in this case.

Interestingly, we found that different adaptive means co-occurred in the evolved populations – independently in different cells or even simultaneously in the same genome. In particular, we saw that evolutionary lines that adapted in *cis* appear to also have had adaptations that are not encoded in the evolving gene, hence pointing to changes that must have occurred in *trans*. Further investigations will reveal which of these solutions, *cis* or *trans*, proves to be more evolutionarily stable - to fully reveal the dynamics of splicing adaptation when cells optimize their gene expression.

## Materials and Methods

### Yeast strains and plasmids

All *S. cerevisiae* strains in this study have the following genetic background: his3Δ1::TEF2-mCherry::URA3::RPS28Ap-YiFP-KAN::NAT; canΔ1::STE2pr-Sp_his5; lypΔ1::STE3pr-LEU2; leu2Δ0; ura3Δ0;

Strains of Y-intron-FP were taken from Yofe *et al^26^* and were introduced with a Kan resistance gene fused 3’ terminally to the YFP. To reconstitute the mutations discovered after lab evolution (rescue strains), we amplified cassettes of Y-i_mut_-FP-KAN and transformed these into the ancestor WT strain, selecting with KAN. Notably, all strains also carry an mCherry-fluorescent protein driven by an independent *TEF2* promoter that was used to normalize cell-to-cell variability for the YFP-Kan expression levels.

### Media

Cultures were grown at 30°C in rich medium (1% bacto-yeast extract, 2% bacto-peptone and 2% dextrose [YPD]). Throughout all experiments, G418 was supplemented to the medium at a concentration of 3mg/ml, which is 10fold higher than the standard.

### Evolution experiments

Lab-evolution experiments were carried out by daily serial dilution for 80 days. Cells were grown on 1.2ml of YPD+G418 at 30°C until reaching stationary phase and then diluted by a factor of 1:120 into fresh media (˜7 generations per dilution, total of ˜560 generations).

### Liquid growth measurements

The cultures were grown at YPD+G418, and optical density (600nm) measurements were taken at 30min intervals. Growth comparisons were performed using 96-well plates, and the growth curve for each strain was obtained by averaging at least 15 wells.

### FACS measurements of YFP-Kan levels

Cells were grown in YPD+G418 at 30°C until they reached the logarithmic growth phase at an optical density of ˜0.4. Then, YFP and mCherry levels were measured for ˜50,000 cells for each culture with flow cytometry. Gating was performed according to side and forward scatters, and YFP levels were normalized with the mCherry signal for each cell individually.

### Quantitative PCR measurements of splicing efficiency

Cultures were grown in YPD+G418 at 30°C until cells reached the logarithmic growth phase at an optical density of ˜0.4. Then, RNA was extracted using MasterPure kit (Epicentre), and were reverse-transcribed to cDNA using random primers. 2μl of cDNA were added to each reaction as template for qPCR using light cycler 480 SYBR I master kit and the LightCycler 480 system (Roche Applied Science), according to the manufacturer’s instructions. For each strain, two qPCRs were performed with three biological repetitions and three technical repeats. A first qPCR was performed targeting the transcript spliced-version with a forward primer complementing the exon-exon junction and a downstream reverse primer. A second reaction targeted the un-spliced version of the transcript with a forward primer complementing the intron and the same reverse primer of the first reaction.

F_exon-exon_ = 5’-CACTACTTTAGGTTATGGTTT-3’

F_intron_ = 5’-CTTCAATTTACTGAATTTGTATG-3’

R_both_ = 5’-GTCTTGTAGTTACCGTCA-3’

Splicing efficiency is reported as the average Cp of the spliced transcript minus the average Cp of the un-splice version.

### mRNA deep sequencing

Cultures were grown in YPD+G418 at 30°C until cells reached the logarithmic growth phase at an optical density of ˜0.4. Cells were then harvested by centrifugation and flash-frozen in liquid nitrogen. RNA was extracted using a modified protocol of nucleospin^®^ 96 RNA kit (Machery-Nagel). Specifically, cell lysis was done in a 96 deep-well plate by adding for each well 450μl of lysis buffer containing 1M sorbitol, 100mM EDTA and 0.45μl lyticase (10IU/μl). The plate was incubated at 30°C for 30min to break cell wall, centrifuged for 10min at 3000rpm, followed by the removal of the supernatant. Then, extraction continued as in the protocol of nucleospin^®^ 96 RNA kit, only using **β**-mercaptoethanol instead of DTT. Poly(A)-selected RNA extracts of size ˜200bps were reverse-transcribed to cDNA using poly(T) primers that were barcoded with a unique molecular identifier (UMI). cDNA was then amplified and sequenced with an Illumina HiSeq 2500.

### Analysis of mRNA deep sequencing

Processing of RNA-seq data was performed as described in Voichek *et al*^43^. Shortly, reads were aligned using Bowtie^44^ (parameters: --best –a –m 2 –strata -5 10) to the genome of *S. Cerevisiae* (R64 from SGD) with an additional chromosome containing the sequence of the YFP-Kan construct. For each sequence, we normalized for PCR bias using UMIs as describe in Kivioja *et al*^45^. Next, reads for each gene end (400bp upstream to 200bp downstream of the ORF’s 3’ end) were summed-up to estimate the gene’s expression level. Genes with coverage lower than 10 reads were excluded. To normalize for differences in coverage among samples, we divided each gene expression by the total read count of each sample and then multiplied by 10^6^. Then, expression ratio was calculated between an evolved/rescue colony to the ancestor and a log2 operation was performed on that ratio. These values were used to compare expression levels of gene groups (ribosomal genes, general stress response genes, splicing machinery genes, intron-containing genes) and of the YFP-Kan mRNA levels as described in the manuscript. When calculating the expression levels of splicing machinery and intron-containing gene groups, the ribosomal and general stress response genes were excluded from the analysis in order to avoid bias from cellular regulation due to changes in physiology and growth rate of the cells.

## Acknowledgments

We thank Martin Mikl for help with the qPCR experiments. We also greatly appreciate comments to the text from Tslil Ast, Orna Dahan, Dvir Schirman, and Avihu Yona, and the entire Pilpel lab for discussions of the project. IF thanks the Azrieli Foundation for an Azrieli PhD Fellowship. This study was supported by the Minerva Foundation who funded the Minerva Center for Live Emulation of Evolution in the Lab.

---

**Supplementary Figure 1.**
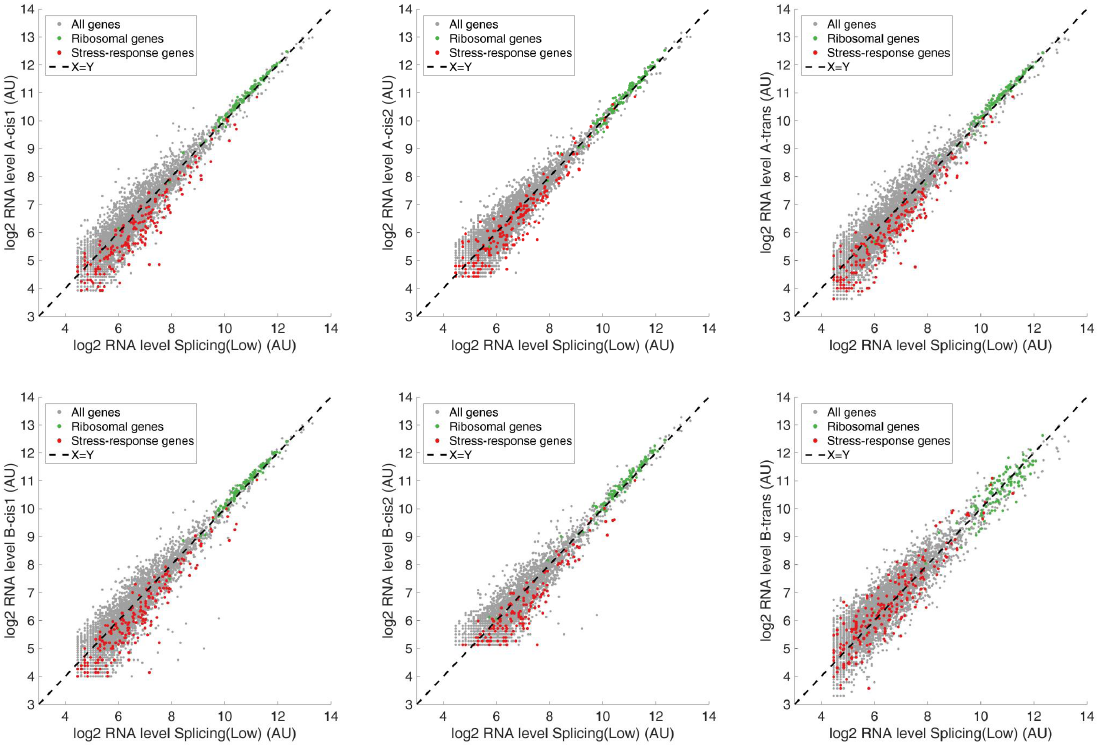
Transcriptome profiling reveals that ribosomal genes were up-regulated and that stress-related genes were down-regulated in 5 out of 6 of the evolved colonies compared to the Splicing^Low^ ancestor.

**Supplementary Figure 2.**
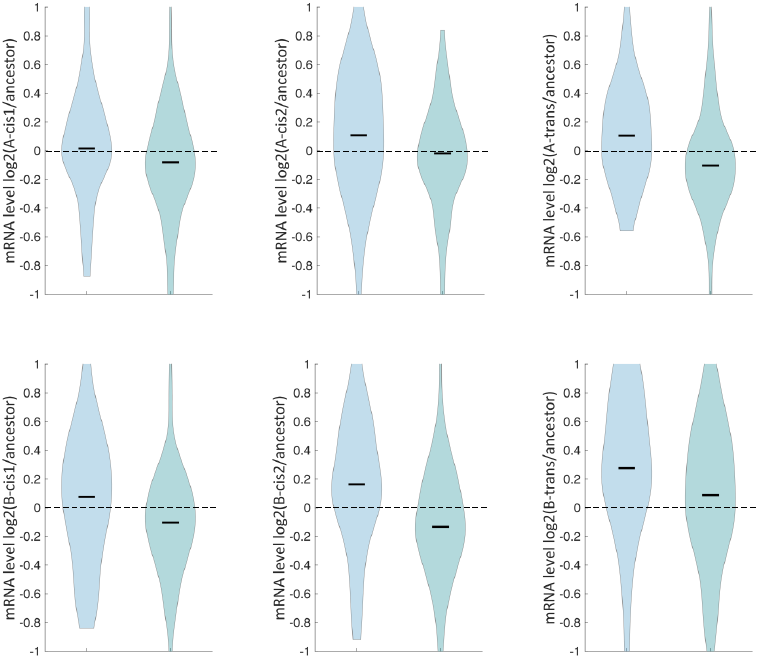
Supply-to-demand ratio in each of the evolved colonies. Folding change ratios in log2 are shown for splicing genes (left) and intron-containing genes (right). Black line represents the median of the distribution.

